# The effects of force constraint during preparatory phase on the explosive force generation of base stealing in baseball

**DOI:** 10.64898/2026.06.10.731238

**Authors:** Kiyohiro Konno, Atsushi Itaya, Tomohiro Kizuka, Seiji Ono

## Abstract

**Background:** Explosive force generation during the initial acceleration phase is critical for successful base stealing in baseball. Preparatory balance control preceding movement onset may facilitate this process by constraining horizontal ground reaction force (GRF) toward a task-specific direction. However, its contribution to ballistic sprint initiation remains unclear.

**Research question:** Does preparatory force constraint influence explosive force generation during base stealing, and when during the preparatory phase is this influence greatest?

**Methods:** Fourteen baseball players performed 3-m maximal sprints simulating base stealing under time-constraint (Time) and self-paced (Self) conditions. GRF around movement onset were recorded. Peak rate of force development (peak RFD) was computed from onset to take-off. A 250-ms window before the onset was divided into 50-ms bins, and mean resultant length (R_len_), which represents the extent of force constraint, of each bin was calculated. Using statistics analyses, Differences between conditions were tested, and the relationship between the interaction (R_len_ × condition) and peak RFD was assessed.

**Results:** The peak RFD was greater under the Self condition than under the Time condition, accompanied by a larger R_len_. Furthermore, Results indicated that the force constraint in the 150–100 ms interval preceding the movement onset most strongly influenced the peak RFD.

**Significance:** These findings demonstrate that temporally organized preparatory force constraint plays a critical role in explosive sprint initiation during base stealing. Identifying the specific preparatory timing linked to superior force production provides novel mechanistic insight into preparatory balance control and may inform targeted training strategies for ballistic athletic movements.

## 1. Introduction

One of the key contributors to increasing scoring opportunities in baseball is a player’s ability to base stealing [1]. A previous study has reported that a one standard deviation increase in the number of stolen base attempts results in 3.65 more games won per season [2]. Although the official distance between adjacent bases is 27.43 m, a runner attempting to steal next base typically takes a lead of approximately 4 m [3]. Thus, the actual running distance is approximately 23 m. Therefore, because the base stealing involves such a short running distance, the ability to exert explosive force, such as rate of force development, is a major factor in successful steals.

For ballistic movements including the base stealing, balance control preceding the movements exerts a significant effect on the performance of the ballistic movements [4–6]. To experimentally examine this preparatory balance control, a classical paradigm manipulates temporal pressure by comparing a time-constrained condition (Time condition), in which participants initiate movement as quickly as possible in response to a cue (e.g., a light stimulus), with a self-paced condition (Self condition). Previous studies employing this paradigm have shown that, relative to the Self condition, the Time condition was associated with insufficient center of pressure (COP) preceding the movements and resulted in deterioration of task performance (e.g., slower onset, reduced shot accuracy) [4,5]. These findings suggest that preparatory balance control plays a critical role in the significant exertion of explosive force.

Force constraint strategy is regarded as a kind of the preparatory balance control. This is high-level neural control strategy organized by central nervous system (CNS) to reduce the degree of freedom in postural control. Although this strategy was originally identified in feedback-based postural responses after movement or external perturbations, subsequent findings suggest that a similar strategy may also be utilized before movement onset through neural activity within the pontomedullary reticular formation (PMRF) [7–10]. As demonstrated by Leonard et al. (2009), preparatory postural adjustments prior to movement generate constrained horizontal ground reaction force (GRF) vectors through the coordinated recruitment of functional muscle synergies [11]. These constrained force vectors shift the center of mass (COM) toward the intended movement direction while maintaining stability within the base of support. Such preparatory organization likely enables the CNS to simplify the control of complex multijoint movements by transforming numerous muscle activations into a limited number of functional force patterns [11,12]. Consequently, the force constraint strategy immediately before movement may contribute to efficient and stable movement execution by optimizing COM displacement, minimizing unnecessary postural variability, and facilitating a smooth transition from posture to voluntary movement. However, the effect of the force constraint strategy preceding base stealing on the explosive force exertion is not directly investigated.

Therefore, aims of this study were twofold. First, to determine whether the force constraint strategy preceding the onset of base stealing influences the generation of explosive force. Second, because previous studies have reported that preparatory force control is gradually modulated as movement onset approaches in short intervals of approximately 50 ms [11,12], to identify the timing of force constraint that most strongly affects the ability to exert maximal force at movement onset. For the aims of this study, we hypothesized that the force constraint parameter under Time condition (i.e., temporal unpredictable) is lesser than that under Self condition (i.e., temporal predictable). Building on this rationale, we additionally hypothesized that the magnitude of the force constraint parameter would predict the ability to exert explosive force at the movement onset.

## 2. Methods

### 2.1. Participants

Fourteen healthy male university baseball players (age: 20.50 ± 1.40 years, height: 172.02 ± 4.55 cm, body weight: 71.93 ± 4.89 kg, years of experience: 13.29 ± 2.09 years, mean ± standard deviation [SD]) participated. Participants with hip, knee, or ankle injury within the past 6 months were excluded. To determine the sample size, a priori Power analysis by G*Power software v3.1.9 (Heinrich Heine University Düsseldorf, Germany, http://www.gpower.hhu.de/) was conducted, and the number of participants was determined based on the results of this power test [13]. Procedures followed the Declaration of Helsinki and were approved by the Research Ethics Committee at the Institute of Health and Sport Sciences, University of Tsukuba. Written informed consent was obtained.

### 2.2. Experimental apparatus and setup

The experimental setup is shown in Figure 1. The participants performed 3 m maximal-effort sprint task simulating base stealing. They wore indoor-rubber shoes and stood with one foot on each of two floor-embedded 3D force plates (9667AA0906, Kistler, size: 900 mm × 600 mm × 100mm) sampled at 1,000 Hz. The right foot position was fixed 0.5 m from the front photoelectric sensors while the left foot was placed naturally on the plate. Two conditions were tested: a light-cue time-constrained condition (Time) and a self-paced condition (Self). In the Time condition, a light-emitting diode (LED) cue positioned at 3 m from away participant signaled sprint start with randomized delays (500–1000 ms) to prevent anticipation. The Self condition was initiated at the participant’s preferred timing. To normalize GRF values at the data processing, they were instructed to maintain bilateral static crouched starting position for 2 seconds (stationary period) at the beginning in all sprint trials. Each condition was performed for 15 trials each (a total of 30 trials). with additional trials collected if stationarity criteria were not met. A 5-min rest separated conditions. The signal of the LED and the force plates were synchronized via an A/D converter (Micro1401, CED) and recorded PC’s application (spike2, Version 5.21, CED).

**Figure 1.**
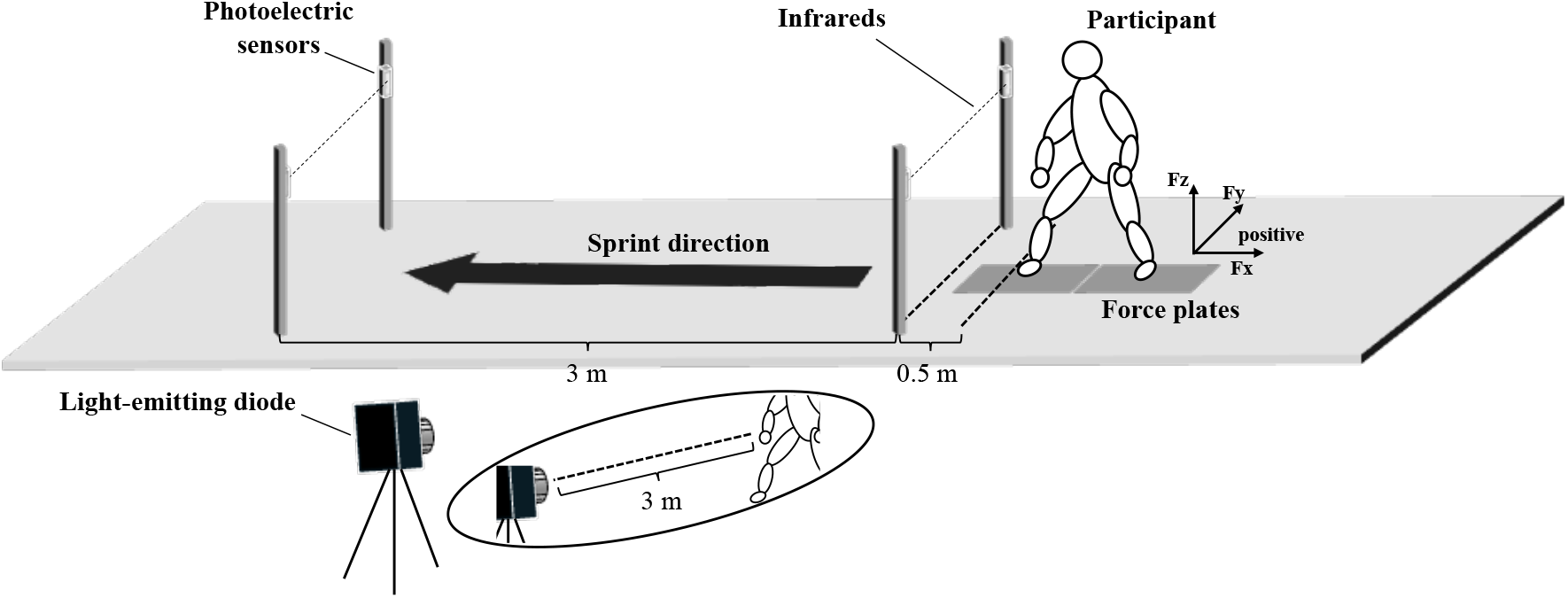
Experimental setup. Participants were instructed to perform 3 m maximal-effort sprint task under conditions where they started at a light-cue (Time condition) or their preferred timing (Self condition). The right foot position of participants was fixed 0.5 m from the front photoelectric sensors while the left foot was placed naturally on the plate. During this experiment, the laboratory coordinate system within force plates was as follows: x-axis = medio-lateral (sprint direction), y-axis = antero-posterior, and z-axis = vertical.

### 2.3. Data processing and analysis

#### 2.3.1. Sprint performance

To quantify the short-distance sprint capability of base stealing, sprint time over the 3 m distance was recorded using a pair of photoelectric sensors placed at the start line and at the goal line. This short distance was chosen because the initial acceleration phase largely determines the outcome of the base stealing [14].

#### 2.3.2. GRF parameters during the propulsion phase

The laboratory coordinate system within force plates was as follows: *x*-axis = medio-lateral (sprint direction), *y*-axis = antero-posterior, and *z*-axis = vertical (Fig 1). Kinetic data were smoothed using a fourth-order Butterworth low-pass digital filter with a cutoff frequency of 15 Hz based on residual analysis [15]. Combined vertical GRF from both plates was used to identify the 0.5 s stationary interval with the smallest SD for body-weight normalization. GRF components were expressed as %BW. Movement onset and take-off were defined when sprint-direction GRF exceeded 5% of its maximum, and when it fell below 5%, respectively. We defined the propulsion phase as the interval from the movement onset to the take-off.

Propulsion duration (pro_dur_) was defined as the time interval from the movement onset to the take-off, and represented the time required to generate sufficient propulsive force to move forward (i.e., sprint-direction). Peak Fx was defined as the maximum value of the sprint-direction GRF observed between the movement onset and the take-off (Fig 2). Furthermore, peak rate of force development (peak RFD) during the propulsion phase was defined as the maximum value of a derivative of the sprint-direction GRF after time-normalization to 100 samples prior to differentiation. The derivatives were computed with respect to cycle fraction. Thus, the derivative values have units of cycle^−1^ (Fig 2).

**Figure 2.**
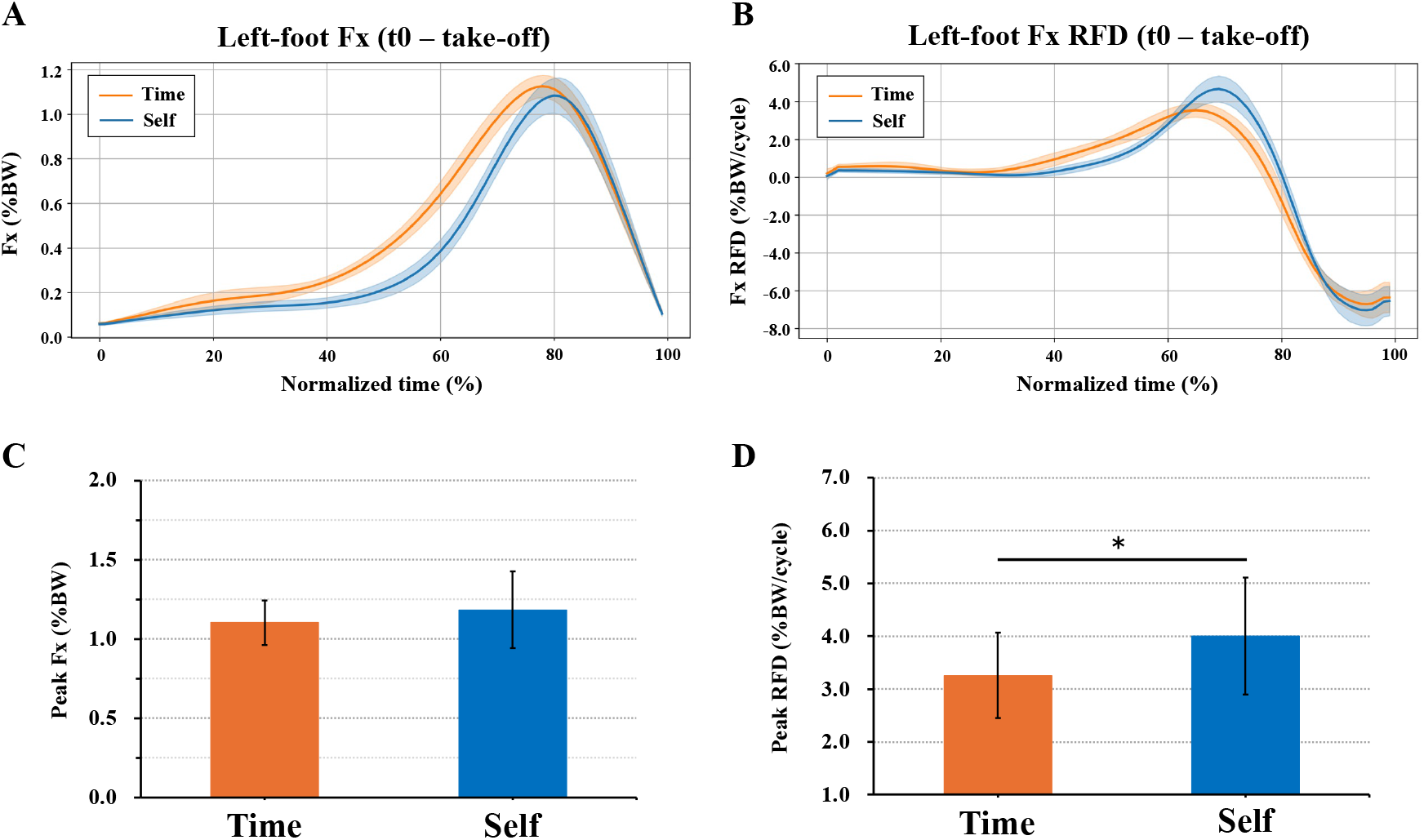
The representative time-normalized data of left-foot Fx and Fx RFD, and the difference of peak Fx and peak RFD between the Time and Self conditions. The time series of Fx were linearly interpolated to 100 samples from the movement onset to take-off (A). The interpolated Fx was then time-differentiated using a central finite-difference to yield the Fx RFD (B). The peak Fx did not differ significantly across conditions (C). In contrast, the peak RFD under the Time condition significantly decreased compared to that under the Self condition (D). The asterisk denoted statistically significant difference across conditions (*p* < 0.05).

#### 2.3.3. Force constraint parameter during preparatory phase

To investigate the temporal organization of force constraint preceding the movement onset (hereafter called the *p*PA phase), the 250 ms preceding the movement onset was subjected to further analysis aimed at quantifying high-level parameters adapted by the CNS. The time lengths were based on well-documented changes in anticipatory postural adjustments preceding voluntary movements [4,16,17]. For each participant and condition, force constraint parameter of each foot was calculated using horizontal force vectors, which refer to the *x*- and the *y*-axes of GRF (Fig 3). Specifically, the 250 ms preceding the movement onset was divided into five 50-ms bins chosen to characterize the modification of the preparation of sprinting movements (e.g., *p*PA1, *p*PA2, etc.). For each 50-ms bin, horizontal angles of each sample in the 50-ms bin were calculated as the arctangent of the GRF of the *x*- and *y*-axes. Moreover, the mean of the horizontal angles in the 50-ms bin was subjected to circular statistical analysis. The mean resultant length (R_len_) represented the indicator of the preparatory force control capability was calculated as:

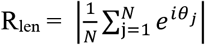

Where *N* is the number of trials each condition, *θ*_*j*_ is the circular mean angle (radians) of trial *j*. Here *e* denotes the base of the natural logarithm, *i* is the imaginary unit, and *e*^*iθ*^ maps an angle to a point on the unit circle via Euler’s formula (i.e., *cosθ + isinθ)*. A standard measure of dispersion for circular data is the R_len_, it takes values between 0 (indicating extreme dispersion) and 1 (indicating perfect constraint around one direction).

**Figure 3.**
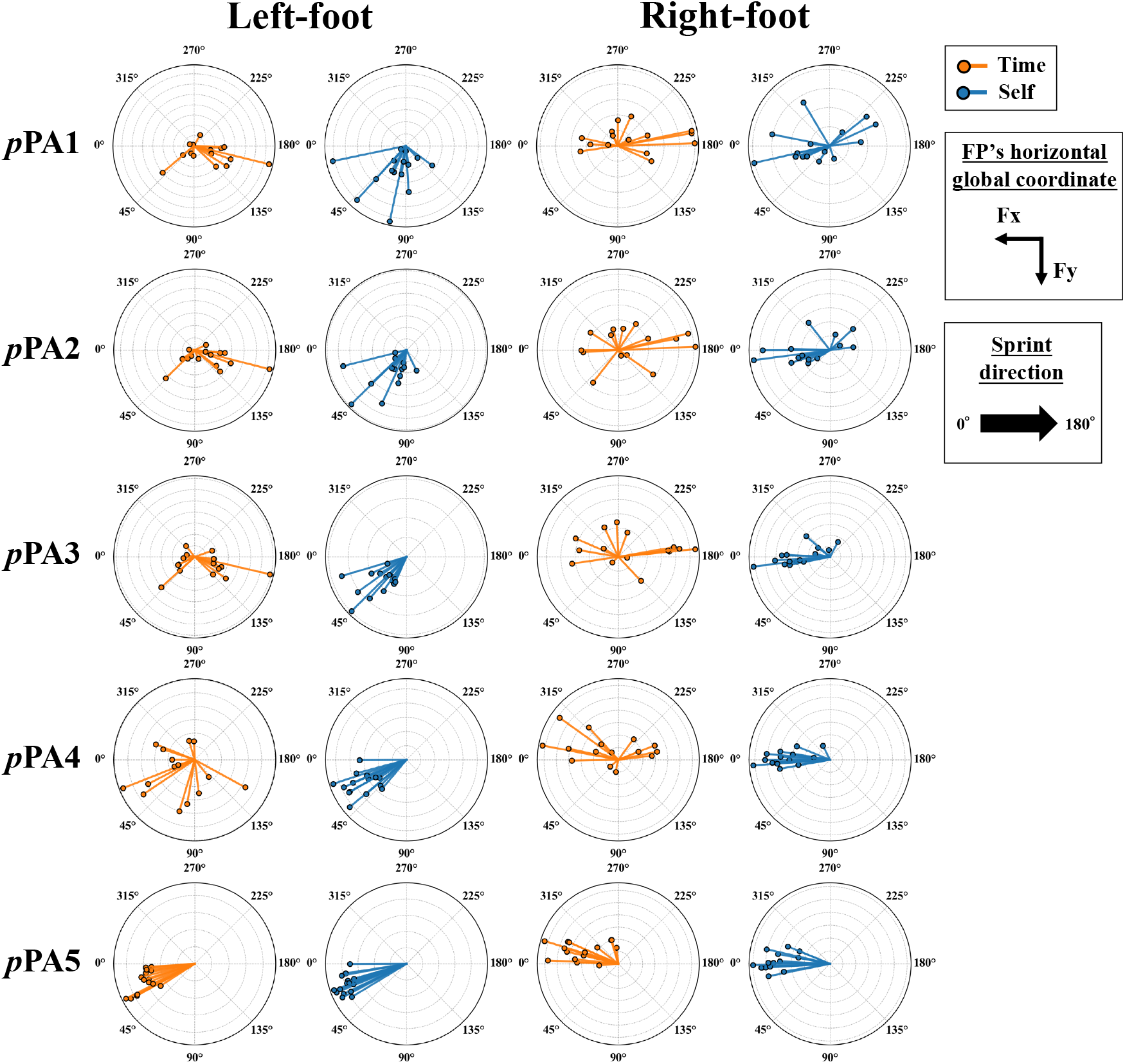
The representative horizontal force constraint during preparatory phase in each trial. The 250-ms interval preceding the movement onset was divided into five 50-ms bins, and defined as *p*PA1, *p*PA2, *p*PA3, *p*PA4, and *p*PA5. For each trial, the composed horizontal vectors (i.e., Fx and Fy) were depicted in each circle, respectively. The composed vectors in each circle biased in a specific direction implies preparatory force constraint, whereas they distributed uniformly implies uncommitted force control (i.e., uncertain timing prediction).

### 2.4. Statistical analysis

Results were reported as mean ± SD. Statistical analyses used IBM SPSS Statistics v30.0 (SPSS Inc., IL, US), and custom scripts in Python v3.13.2. Normality was assessed with Shapiro–Wilk test, and sphericity violations were corrected with Greenhouse–Geisser. Sprint time differences were tested with Wilcoxon signed-rank tests while paired *t*-tests compared GRF parameters. For the force constraint parameter, left-foot data met normality and were analyzed with two-way (i.e., bins × conditions) repeated-measures ANOVA (tw-rmANOVA). Right-foot data were analyzed with aligned rank transform (ART) tw-rmANOVA because it violated normality [18]. Relationships between predictors and outcomes were examined with linear mixed-effects (LME) models including fixed effects for condition, predictor, and their interaction, and a random intercept for participant. Models were estimated by maximum likelihood. Two models (full model: with condition × predictor interaction, reduced model: without the interaction) were compared by likelihood-ratio (LR) tests to justify reporting simple slopes. LME model residual normality and homoscedasticity were checked with Shapiro–Wilk and Breusch–Pagan tests. Effect sizes were reported for all analyses: Cohen’s *d* for paired *t*-test (small ≈ 0.2, medium ≈ 0.5, large ≈ 0.8), *r* for Wilcoxon signed-rank test (small ≈ 0.1, medium ≈ 0.3, large ≈ 0.5), *η*_*p*_^*2*^ for tw-rmANOVA (small ≈ 0.01, medium ≈ 0.06, large ≈ 0.14) [19]. The significance level for all the statistical analyses was set at 0.05.

## 3. Results

### 3.1. Sprint performance

The result of Wilcoxon signed-rank test for the sprint time showed that there was no significant difference between the Time and the Self conditions (Time, 0.69 ± 0.03 s, Self, 0.68 ± 0.03 s, *Z*_*14*_ = - 1.35, *p* = 0.18, *r* = 0.25).

### 3.2. GRF parameters during the propulsion phase

Paired *t*-tests were performed for the pro_dur_, the peak Fx, and the peak RFD. For the pro_dur_, there was no significant difference between the Time and the Self conditions (Time, 0.32 ± 0.47 s, Self, 0.34 ± 0.57 s, *t*_*13*_ = -1.14, *p* = 0.28, *d* = -0.30). Similarly, the peak Fx did not differ significantly across conditions (Time, 1.10 ± 0.14 %BW, Self, 1.18 ± 0.24 %BW, *t*_*13*_ = -1.57, *p* = 0.14, *d* = -0.42, Fig 2). In contrast, the peak RFD showed a significant difference, with values larger in the Self condition than in the Time condition (Time, 3.26 ± 0.81 %BW/cycle, Self, 4.00 ± 1.11 %BW/cycle, *t*_*13*_ = -4.12, *p* = 1.00 × 10^−3^, *d* = -1.10, Fig 2).

### 3.3. Force constraint parameters during preparatory phase

For the left foot, there was a significant main effect of phase (*p*PA1, 0.58 ± 0.03 a.u., *p*PA2, 0.62 ± 0.04, *p*PA3, 0.66 ± 0.03, *p*PA4, 0.71 ± 0.03, *p*PA5, 0.84 ± 0.03 a.u., *F*_*(4,52)*_ = 13.89, *p* = 1.40 × 10^−4^, *η*_*p*_^*2*^ = 0.52, Fig 4) as well as a significant main effect of condition (Time, 0.58 ± 0.05, Self, 0.79 ± 0.03 a.u., *F*_*(1,13)*_ = 12.50, *p* = 4.00 × 10^−3^, *η*_*p*_^*2*^ = 0.49, Fig 4). No significant interaction between phase and condition was found (*F*_*(4,52)*_ = 2.61, *p* = 0.01, *η*_*p*_^*2*^ = 0.17). In addition, the result on the right-foot R_len_ was shown that a main effect of phase was statistically significant (*p*PA1, 0.49 ± 0.27, *p*PA2, 0.56 ± 0.24, *p*PA3, 0.58 ± 0.28, *p*PA4, 0.66 ± 0.27, *p*PA5, 0.87 ± 0.13 a.u., *F*_*(4,117)*_ = 17.19, *p* = 4.10 × 10^−11^, *η*_*p*_^*2*^ = 0.37, Fig 4), and the other main effect was also significant (Time, 0.59 ± 0.29, Self, 0.67 ± 0.25 a.u., *F*_*(1,117)*_ = 5.52 *p* = 0.02, *η*_*p*_^*2*^ = 0.05, Fig 4). No significant interaction between phase and condition was found (*F*_*(4,117)*_ = 0.69, *p* = 0.60, *η*_*p*_^*2*^ = 0.02, Fig 4).

**Figure 4.**
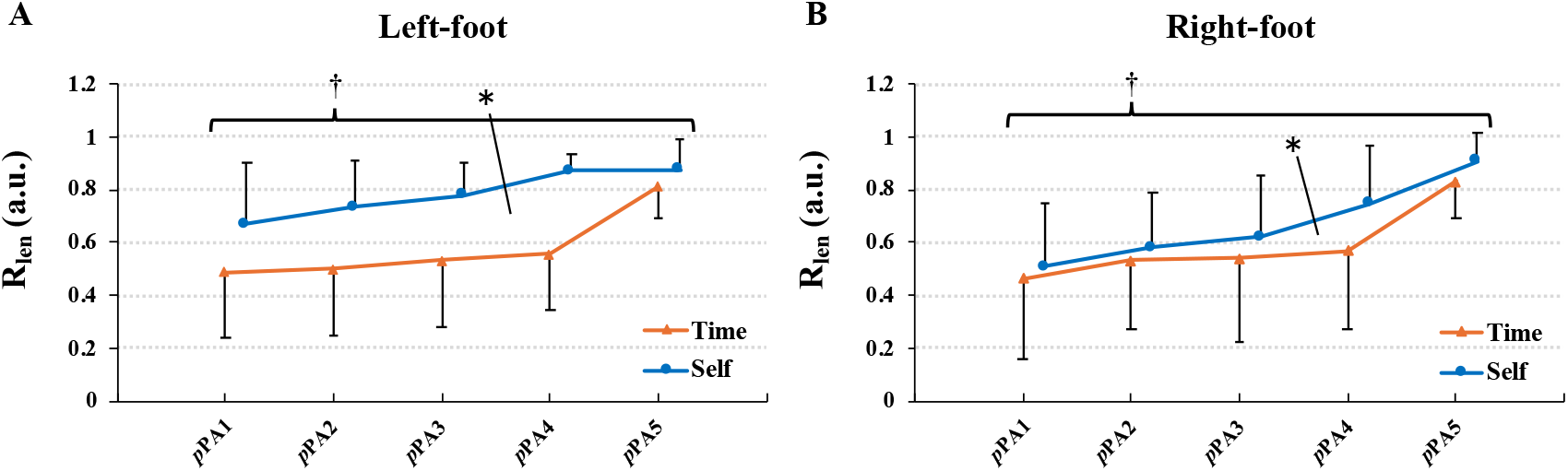
The differences of the force constraint parameters in each bin, condition, and foot. The mean resultant length (R_len_) was calculated by circular statistical analysis. For each foot, there were significant main effects of bins and conditions (A: Left-foot R_len_, B: Right-foot R_len_). The asterisks and daggers denote statistically significant difference across conditions and bins, respectively (*p* < 0.05).

### 3.4. Relationships between force constraint parameter in each bin and peak RFD

All result of LR tests for LME models and fitting were presented at Table 1.

**Table 1.**
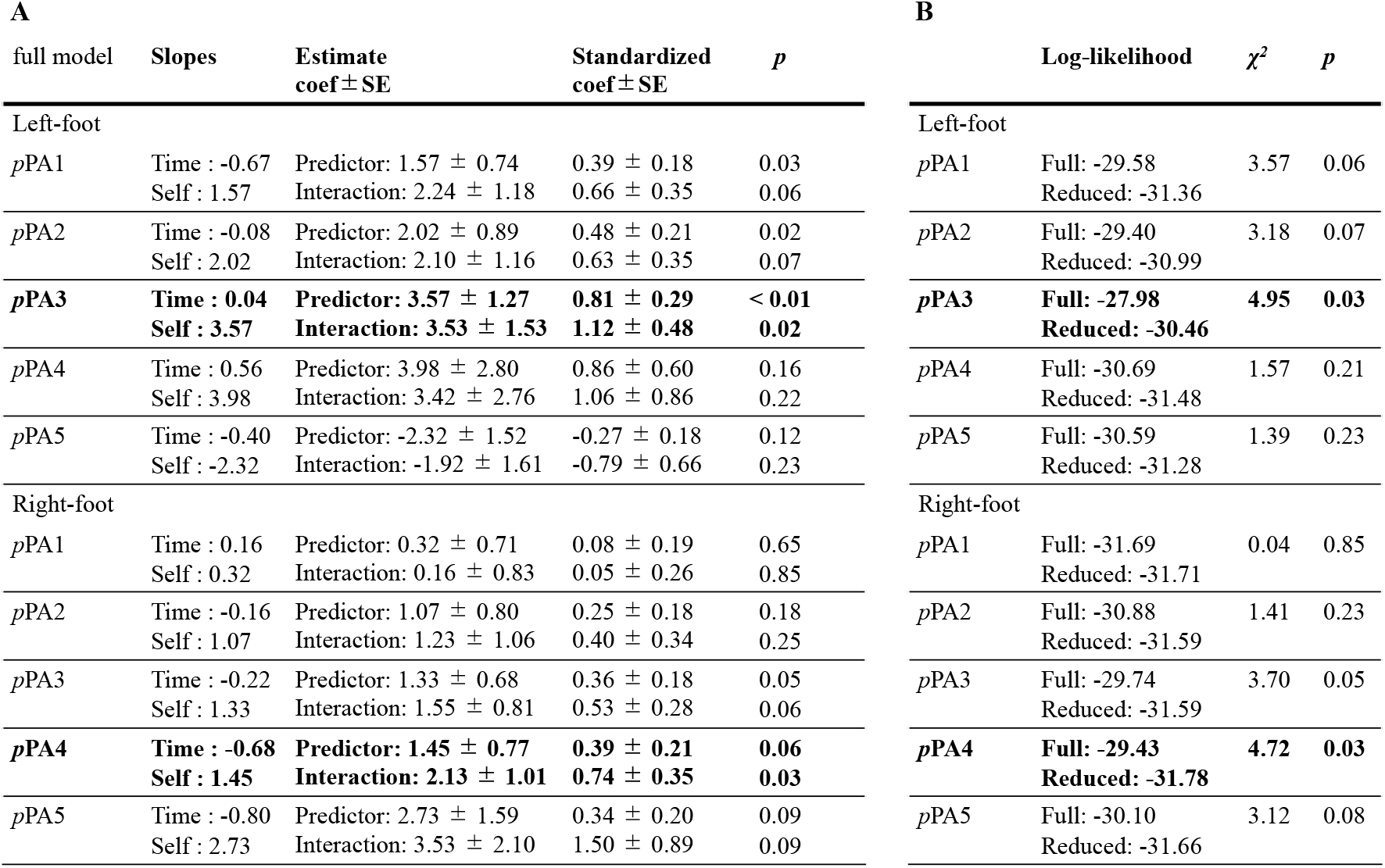
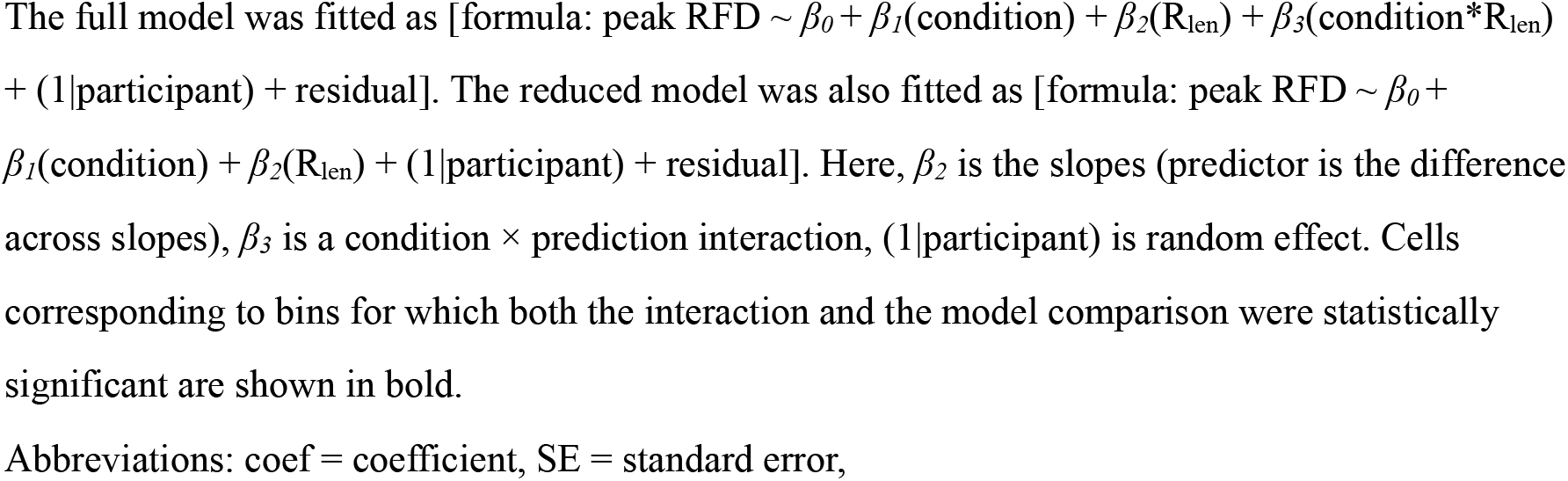
Summary of full model fits and LME models with/without interaction comparisons.

LME model comparisons between the full and reduced models showed that including the condition × R_len_ interaction improved fit for left-foot *p*PA3 and right-foot *p*PA4 (*p*PA3, *χ*^*2*^ = 4.95, *p* = 0.03, *p*PA4, *χ*^*2*^ = 4.72, *p* = 0.03). In the full model, the slope for the Time condition (relative to the Self condition) significantly differed (left-foot *p*PA3; Time slope = 0.04, Self slope = 3.57, *p* = 0.02, *β* = 1.12, right-foot *p*PA4; Time slope = -0.68, Self slope = 1.46, *p* = 0.03, *β* = 0.74, Fig 5). These standardized coefficients (*β*) indicate that a one SD increase in the predictor is associated with increases in peak RFD of 1.12 and 0.74 SDs for left-foot *p*PA3 and right-foot *p*PA4, respectively.

**Figure 5.**
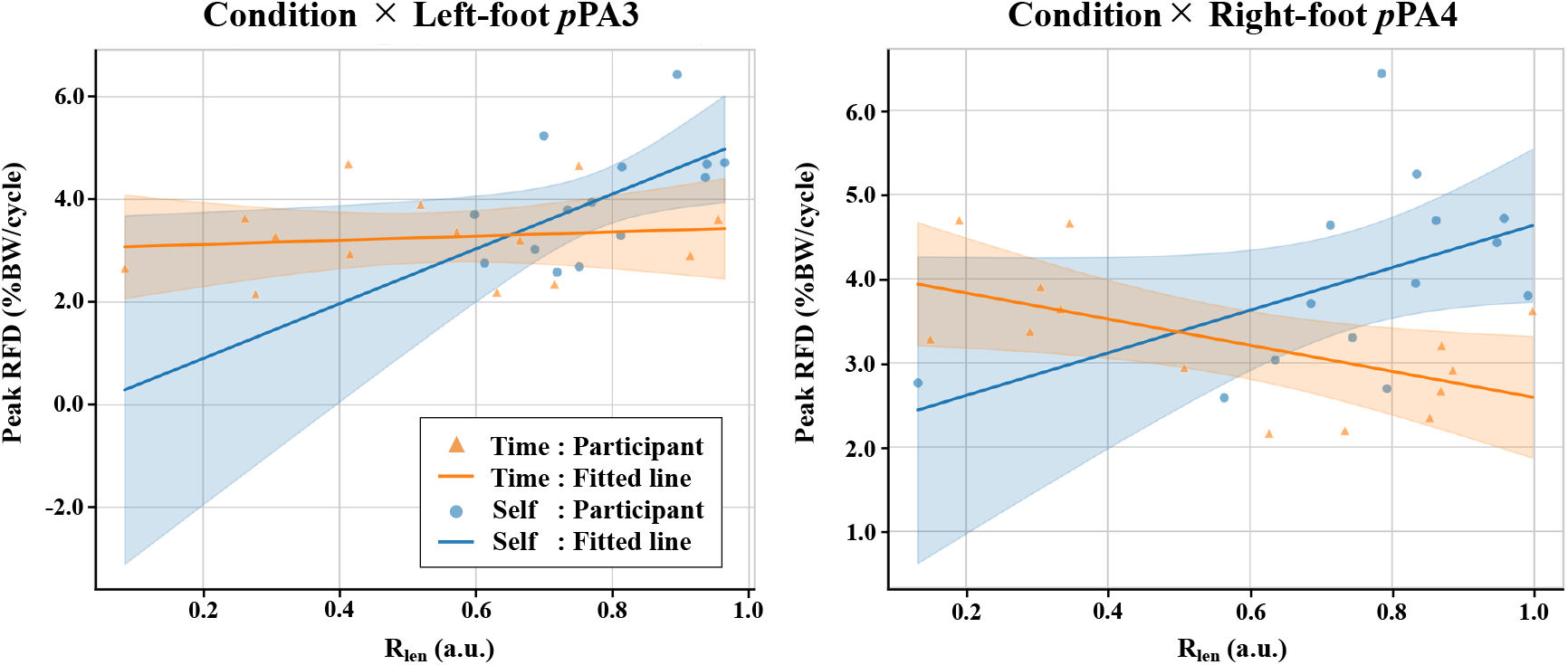
Relationship between mean resultant length (R_len_) and peak rate of force development (Peak RFD). Scatter points show participant-level representative values. Orange triangles denote the Time condition, and blue circles denote the Self condition. Solid lines are condition-specific linear fits with shaded 95% confidence bands. The fitted lines illustrate the divergent slopes for Time (orange) and Self (blue).

## 4. Discussion

The aims of this study were to determine whether the force constraint strategy preceding the onset of base stealing influences the generation of explosive force, and to identify the timing of force constraint that most strongly affects the ability to produce maximal force at movement onset. To this end, we employed the classical paradigm and compared the indicators reflecting the ability of explosive force exertion and force constraint strategy preceding the movement onset between the Time and Self conditions.

In this study, the result showed that the sprint time was not the significant difference between the Time and Self conditions. We consider that a sprint parameter, such as the sprint time in this study, is influenced by various factors. In particular, previous studies have reported the importance of exerting greater explosive force through hip extension and ankle adduction during acceleration phase (i.e., immediately after the movement onset) [14,20,21]. That is, these findings indicate that the sprint time can be influenced by the body’s kinetics and kinematics immediately after the movement onset. Alternatively, the distance used in this study may have been short for differences in sprint performance associated with temporal pressure to be reflected in overall sprint time [14]. Therefore, the sprint time defined in this study may not have shown a statistically significant difference between the Time and Self conditions. However, the peak RFD differed significantly between the Time and Self conditions, while other GRF parameters showed no statistical change. The RFD is a well-established determinant of early sprint acceleration, with larger RFD values linked to superior short-distance performance [22,23]. Thus, the result in this study indicates that imposing time constraints compromises the preparatory force control to generate explosive force at the movement onset, even when maximum force and duration remain comparable.

The above mentioned seems to be substantiated by the results that the differences between the Time and Self conditions for the force constraint parameters preceding the propulsion phase. In line with our hypotheses presented in the Introduction, significant differences of force constraint parameter in each foot were observed between the Time and Self conditions. These results were consistent with previous findings that the onset of preparatory balance control under time pressure condition was later than that under self-paced condition [4,5,24]. These findings indicate that temporal pressure modifies force organization pre-planned by CNS, reducing the sufficiency of preparation and thereby disturbing the generation of explosive force at the movement onset. Significantly, during the preparatory phase, we found the timing of force constraints that most strongly affects the ability to produce maximal force at movement onset (i.e., *p*PA3 of the left-foot R_len_ and *p*PA4 of the right-foot R_len_). This is because, to our knowledge, no previous study has identified which specific preparatory phase exerts the greatest influence on performance output. Based on several studies examining preparatory balance control, the organization depends on individual’s previous experience (e.g., beginner and expert) and specific tasks (e.g., focal and whole-body movement) [25,26]. Because force constraint preceding the movement onset is shaped by sensorimotor experience and practice, it undergoes adaptive modification through learning [27,28]. Therefore, although these findings may be specific to the base stealing in baseball and to participants possessing the particular task-relevant experience represented in our sample (i.e., trained players), they nonetheless provide important mechanistic insight into how experience-dependent preparatory force strategies are temporally organized and scaled, with direct implications for targeted training interventions and for broader theories of preparatory motor control.

## Conclusion

This study investigated the effect of force constraint strategy preceding the movement onset on sprint performance in base stealing. These findings demonstrate that temporally organized preparatory force constraint plays a critical role in explosive sprint initiation during base stealing. Identifying the specific preparatory timing (i.e., left-foot *p*PA3 and right-foot *p*PA4) linked to superior force production provides novel mechanistic insight into preparatory balance control and may inform targeted training strategies for ballistic athletic movements.

## Acknowledgements

We appreciate Takato Mori’s, as an examiner, contributions to participant recruitment and to the successful completion of the data collection.

